# A space and time-efficient index for the compacted colored de Bruijn graph

**DOI:** 10.1101/191874

**Authors:** Fatemeh Almodaresi, Hirak Sarkar, Rob Patro

**Author notes:** The authors contributed equally to this work (Fatemeh Almodaresi, Hirak Sarkar).

## Abstract

We present a novel data structure for representing and indexing the compacted colored de Bruijn graph, which allows for efficient pattern matching and retrieval of the reference information associated with each k-mer. As the popularity of the de Bruijn graph as an index has increased over the past few years, so have the number of proposed representations of this structure. Existing structures typically fall into two categories; those that are hashing-based and provide very fast access to the underlying k-mer information, and those that are space-frugal and provide asymptotically efficient but practically slower pattern search.

Our representation achieves a compromise between these two extremes. By building upon minimum perfect hashing, carefully organizing our data structure, and making use of succinct representations where applicable, our data structure provides practically fast k-mer lookup while greatly reducing the space compared to traditional hashing-based implementations. Further, we describe a sampling scheme built on the same underlying representation, which provides the ability to trade off k-mer query speed for a reduction in the de Bruijn graph index size. We believe this representation strikes a desirable balance between speed and space usage, and it will allow for fast search on large reference sequences.

Pufferfish is developed in C++11, is open source (GPL v3), and is available at https://github.com/COMBINE-lab/Pufferfish. The scripts used to generate the results in this manuscript are available at https://github.com/COMBINE-lab/pufferfish_experiments.

## 1 Introduction

Motivated by the tremendous growth in the availability and affordability of high-throughput genomic, metagenomic and transcriptomic sequencing data, the past decade has seen a large body of work focused on developing data structures and algorithms for efficiently querying large texts (e.g. genomes or collections of genomes)^1,2,3,4,5,6,7,8,9^. While numerous approaches have been proposed, many fall into one of two categories — those based on indexing fixed-length pattern occurrences (i.e., k-mers) in the reference sequences ^1,4,7^(most commonly using hashing), and those based on building full-text indices such as the suffix array or FM-index over the references ^2,3,5,6,8,9^.

Recently, there have been efforts to extend both categories of approaches from the indexing of linear reference genomes to the indexing of different types of sequence graphs^10^, with various tradeoffs in the resulting space and time efficiency. On the full-text index side, examples include approaches such as those of Maciuca et al. ^11^and Beller and Ohlebusch^12^ which encode the underlying graph using variants of the BWT, and the approach of Sirén^13^, which indexes paths in the variation graph (again making use of a modified BWT). There have also been recent approaches based on k-mer-indices that adopt graphs as the underlying representation of the text being searched. Examples of such tools include genomeMapper ^14^, BGREAT^15^, kallisto^16^ and deBGA ^17^.

Rather than general variation graphs, we focus in this manuscript on the de Bruijn graph. The de Bruijn graph is a widely-adopted structure for genome and transcriptome assembly,^18,19,20^. However, the compacted variant of the de Bruijn graph has recently been gaining increasing attention both as an indexing data structure—for use in read alignment^17^ and pseudoalignment^16^—as well as a structure for the analysis of variation (among multiple genomes)^21^. The compacted de Bruijn graph ^22,23,24^ is particularly attractive for representing and indexing repetitive sequences, since exactly repeated sequences of length at least *k* are represented only once in the set of unique, non-branching paths. As has been demonstrated by Liu et al.^17^, this considerably speeds up alignment to repeat-heavy genomes (e.g., the human genome) as well as to collections of related genomes.

However, the query speed of existing compacted de Bruijn graph indices comes at a considerable cost in index size and memory usage. Specifically, the need to build a hash table over the k-mers appearing in the de Bruijn graph unipaths requires a large amount of memory, even for genomes of moderate size. Typically, these hash functions map each k-mer (requiring at least 8 bytes) to the unipath in which it occurs (typically 4 or 8 bytes) and the offset where the k-mer appears in this unipath (again, typically 4 or 8 bytes). A number of other data structures are also required, but, most of the time, this hash table dominates the overall index size. For example, an index of the human genome constructed in such a manner (i.e., by deBGA or kallisto) may require 40—100GB of RAM (see Table 2). This already exceeds the memory requirements of moderate servers (e.g., those with 32G or 64G of RAM), and these requirements quickly become untenable with larger genomes or collections of genomes.

**Fig. 1:**
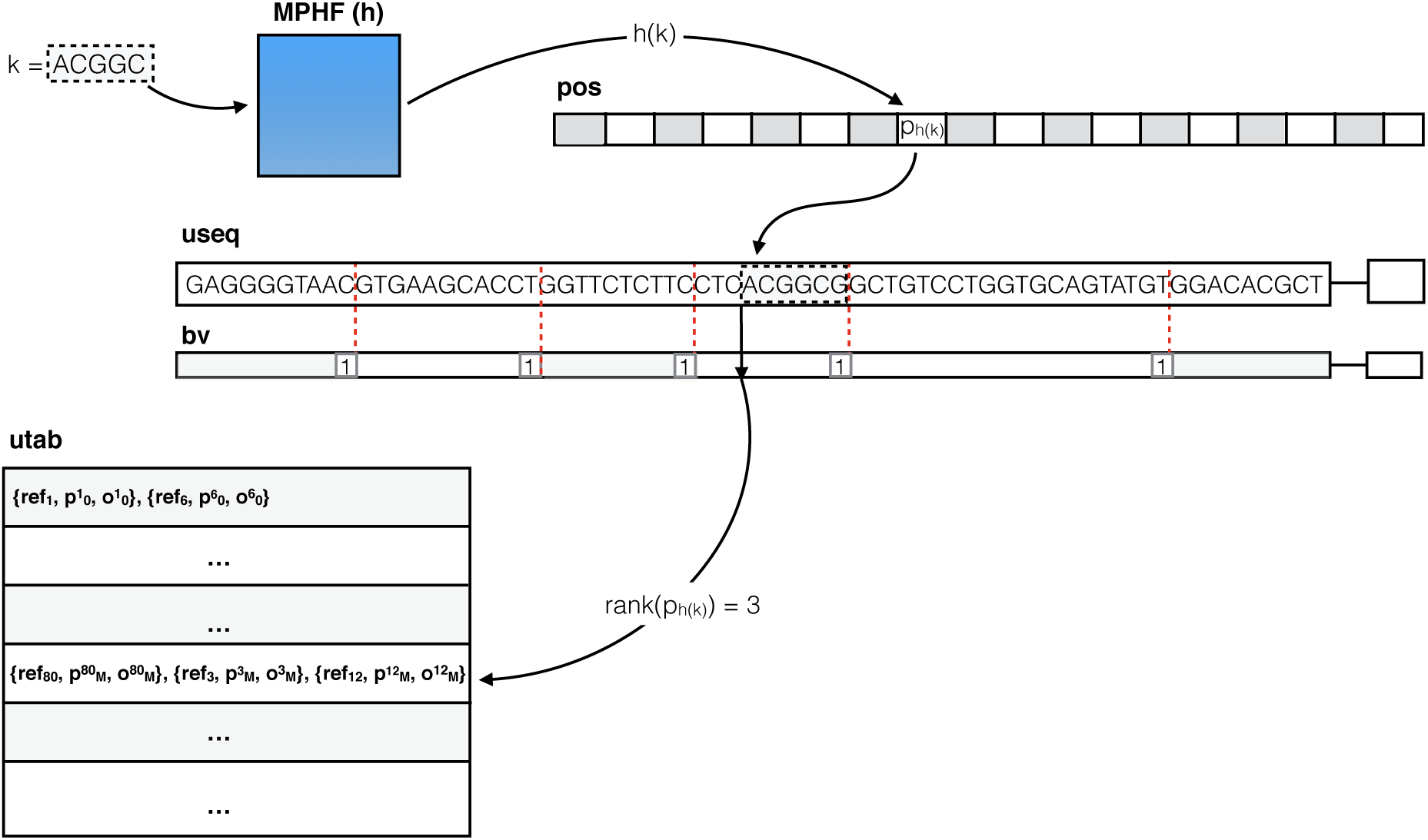
An illustration of searching for a particular k-mer in the *dense* Pufferfish index. The minimum perfect hash yields the index, 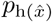 in the pos vector where the k-mer appears in the unipath array. The k-mer is validated against the sequence recorded at this position in `useq` (and, in this case, it matches). A rank operation on 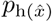 is performed in the boundary vector (`bv`), which yields the corresponding unipath-level information in the unipath table (`utab`). If desired, the relative position of the k-mer within the unipath can be retrieved with an extra select and rank operation.

**Table 2:**
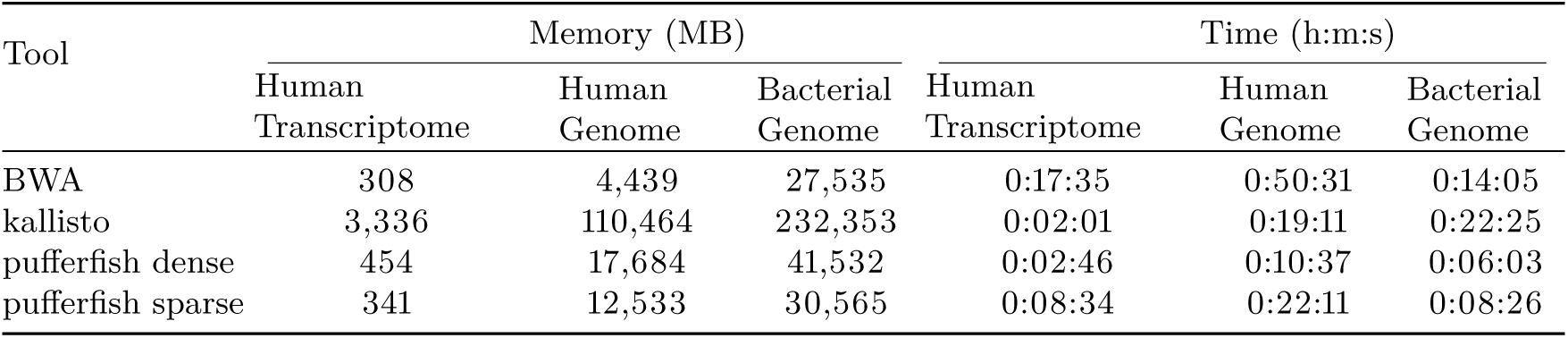
The time and memory required load the index and query all k-mers in reads of the input FASTQ files for different datasets.

## 2 Methods

We present Pufferfish, a software tool implementing a novel indexing data structure for the compacted de Bruijn graph and the colored compacted de Bruijn graph. We focus on making the compacted de Bruijn graph index practical in terms of disk and memory resources for genomic and metagenomic data while maintaining very fast query speeds over the index. While we are conscious of memory usage, we don’t aim to build the smallest possible index. Furthermore, we introduce two different variants of our index, the *dense* and *sparse* Pufferfish indices. Similar to the FM-index ^25^, in the *sparse* Pufferfish index, there is a sampling factor that can be tuned to trade off search speed for index size. The dense index is, in a sense, just a variant of the sparse index tuned for maximum speed (and, hence, taking maximum space). However, as we believe the dense index will be a popular choice, we implement a few optimizations and describe the structures separately.

*Pre-processing* We assume as input to Pufferfish the compacted de Bruijn graph on the reference or set of references to be indexed. The Pufferfish software itself accepts as input a graphical fragment assembly (GFA) format^*^ file that describes the compacted de Bruijn graph. Specifically, this file encodes the unipaths (i.e., non-branching paths) of the compacted de Bruijn graph as “segments” and the mapping between these unipaths and the original reference sequences as “paths”. Each path corresponds to an input reference sequence (e.g., a genome), and is spelled out by an ordered set of unipath IDs and the orientation with which these unipaths map to the reference, so that each unipath has an overlap of *k* – 1 with its following unipath in the path (either in the forward or reverse-complement direction).

GFA is an evolving standard that is meant to be a common format used by tools dealing with graphical representations of genomes or collections of genomes. We note that there are a number of software tools for building the compacted de Bruijn graph directly (i.e., without first building the un-compacted de Bruijn graph). We adopt TwoPaCo ^21^, which employs a time and memory-efficient parallel algorithm for directly constructing the compacted de Bruijn graph, and whose output can be easily converted into GFA format. We note that, due to a technical detail concerning how TwoPaCo constructs the compacted de Bruijn graph and the GFA file, the output cannot be directly used by Pufferfish. Therefore, the current workflow of Pufferfish includes a GFA-to-GFA converter that prepares the TwoPaCo-generated GFA file for indexing by Pufferfish^**^.

### The dense Pufferfish index

Here we describe the basic (i.e., dense) Pufferfish index. The index consists of 6 components (one of which is optional) as described below, and the overall structure is similar to what is explained by Liu et al.^17^:

useq: The unipath sequence array (`useq`) consists of the (2-bit encoded) sequence of all unipaths of the compacted de Bruijn graph packed together into a single array. Typically, the size of this structure is close to (or smaller than) the size of the 2-bit encoded reference sequence, since redundant sequences are represented only once in this structure. We note that the unipath array contains the sequence of every valid k-mer, as well as that of potentially invalid k-mers (those which span unipath boundaries in the packed array, as the sequences in the array follow each other without any delimiters or gaps.). We denote by *L*_*s*_ the total length (in nucleotides) of the unipath array.
bv: The boundary vector (`bv`) is a bit-vector of length *L*_*s*_. The bits of this vector are in one-to-one correspondence with the nucleotides of the unipath array, and the boundary vector contains a one at each nucleotide corresponding to the end of a unipath in `useq`, and a zero everywhere else. We can retrieve the index of each unipath in `useq` using the rank operation on `bv`. `rank`(`bv`[*i*]) returns the number of 1s in bv before the current index, *i*, or, in other words, the index of the current unipath. This can be used to get reference information for the current unipath from utab, which is explained below. We note that bv is typically *very* sparse, and so can likely be compressed (using e.g., RRR^26^ or Elias-Fano encoding), though we have not explored this yet.
h: The minimum perfect hash function (h) maps every *valid* k-mer in the unipath array (i.e., all k-mers not spanning unipath boundaries) to a unique number in [0, *N*), where *N* is the number of distinct valid k-mers in `useq`. We make use of the highly-scalable MPHF construction algorithm of Limasset et al.^27^. We also note that we build the MPHF on the canonicalized version of each k-mer.
pos: The position vector (`pos`) stores, for each valid k-mer *x*, the position where this k-mer occurs in `useq`. Specifically, for k-mer *x*, let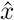 be the reverse complement of *x* and let 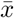 be the canonical form of *x* (the lexicographically smaller of *x* and 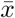). Then `pos` 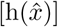 contains the starting position of *x* in `useq` such that `useq` 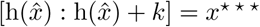
utab: The unipath table `utab` stores, for each unipath appearing in `useq`, the reference sequences (including reference ID, offset and orientation) where this unipath appears in the reference. This is similar to a “posting list” in traditional inverted indices, where all occurrences of the item (in this case, an entire compacted de Bruijn graph unipath) are listed. The order of the unipaths in utab is the same as their order in `useq`, allowing the information for a unipath to be accessed via a simple rank operation on `bv`.
eqtab: *Optionally*, an equivalence class table that records, for each unipath, the set of reference sequences where this unipath appears. Pre-computation and storage of these equivalence classes can speed up certain algorithms (e.g., pseudoalignment^16^).W)

These structures allow us to index every k-mer in the compacted de Bruijn graph efficiently, and to recall, on demand, all of the reference loci where a given k-mer occurs. We note here that the k-mers of the compacted de Bruijn graph constitute only a subset of the k-mers in `useq`. We refer to all k-mers in `useq` that do not span the boundary between two unipaths as *valid* k-mers; these are in one-to-one correspondence with the k-mers of the compacted de Bruijn graph.

**k-mer query in the dense Pufferfish index** By using a minimum perfect hash function (MPHF), h, to index the *valid* k-mers, we avoid the typically large memory burden associated with standard hashing approaches. Instead, the identity of the hashed keys is encoded implicitly in `useq`. Given a k-mer *x*, we can check for its existence and location in the following way. We first compute *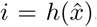*, the index assigned to k-mer *x* by *h*. If *i ≥ N*, then we immediately know that *x* is not a valid k-mer. Otherwise, we retrieve the position *p*_*i*_ stored in pos[*i*]. Finally, we check if the encoded string `useq`[*p*_*i*_ : *p*_*i*_ + *k*] is identical to *x* (or 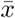). If so, we have found the unipath location of this k-mer. Otherwise, *x* is not a valid k-mer.

Given *p*_*i*_, we can retrieve the reference positions by computing *r*_*pi*_ = `rank`(`bv`[*pi*]), which provides an index into `utab` that is associated with the appropriate unipath. This provides all of the reference sequences, offsets and orientations where this unipath appears. We compute the offset of k-mer *x* in the unipath as *o*_*i*_ = *p*_*i*_ `select` (*r*_*pi*_), where `select` (*r*_*pi*_) returns the start position of the unipath in utab. This allows us to easily project this k-mer’s position onto each reference sequence where it appears. We note that querying a k-mer in the Pufferfish index is an asymptotically constant-time operation, and that the reference loci for a k-mer *x* can be retrieved in 𝒪 (`occ`(*x*)) time, where occ(*x*) is the number of occurrences of *x* in the reference.

### The sparse Pufferfish index

The Pufferfish index, as described above, is relatively memory-efficient. Yet, what is typically the biggest component, the pos vector, can still grow rather large. This is because it requires [lg(*L*_*s*_)] bits for each of the *N* valid k-mers in `useq`. However, at the cost of a slight increase in the practical (though not asymptotic) complexity of lookup, the size of this structure can be reduced considerably. To see how, we first make the following observation:

**Observation 1** *In the compacted de Bruijn graph (and hence, in `useq`), each valid k-mer occurs exactly once (k-mers occuring between unipath boundaries are not considered). Hence, any valid k-mer in the compacted de Bruijn graph is either a complex k-mer (i.e., it has an in or out degree greater than 1), is a terminal k-mer (i.e., it appears at the beginning or end of some input reference sequence), or it has a unique predecessor and / or successor in the orientation defined by the unipath.*

We can exploit this observation in Pufferfish to allow *sampling* of the k-mer positions. That is, rather than storing the position of each k-mer in the unipath array, we store the position only for some subset of k-mers, where the rate of sampling is given by a user-defined parameter *s*. For those k-mers that are not sampled, we store, instead, three pieces of information; the extension that must be applied to move toward the closest k-mer at a sampled position (the `QueryExt` vector), whether or not the corresponding k-mer in `useq` is canonical (the `isCanon` vector), and whether the extension to reach the nearest sampled position should be applied by moving to the right or the left (the `Direction` vector). The `QueryExt` vector encodes the extensions in a 3-bit format so that variable-length extensions can be encoded, though every entry in this vector is reserved to take the same amount of space (3 times the maximum extension length, *e*). The `isCanon` vector is set to 1 whenever the corresponding k-mer appears in `useq` in the canonical orientation, and is set to 0 otherwise. The `Direction` vector is set to 1 whenever the corresponding, non-sampled, k-mer should be extened to the right, and it is set to 0 when the corresponding k-mer should be extended to the left. We additionally store an extra bit vector with the same size as `useq` (the `isSamp` vector) that is set to 1 for any k-mer whose position is sampled and 0 for all other k-mers.

This idea of sampling the positions for the k-mers is similar to the idea of sampling the suffix array positions that is employed in the FM-index ^25^, and the idea of walking to the closest sampled position to verify a k-mer occurs is closely related to the shallow forest covering idea described by Belazzougui, Djamal and Gagie, Travis and Mäkinen, Veli and Previtali, Marco^28^ for verifying membership of a k-mer in their fully-dynamic variant of the de Bruijn graph. This scheme allows us to trade off query time for index space, to allow the Pufferfish index to better scale to large genomes or collections of genomes.

**k-mer query in the sparse Pufferfish index** k-mer query in the sparse Pufferfish index is the same as that in the dense index, except for the first step — determining the position of the k-mer *x* in `useq`. When we query the MPHF with *x* to obtain *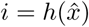*, there are three possible results.

1. In the first case, if *i ≥ N*, this implies, just as in the dense case, that *x* is not a valid k-mer.

2. In the second case, if *i < N* and `isSamp`[*i*] = 1, this implies that we have explicitly stored the position for this k-mer. In this which case we can retrieve that position as *p*_*i*_ = `pos`[`rank`(`isSamp`[*i*])] and proceed as before in the dense case to validate *x* and retrieve its reference positions.

3. In the third case, if *i < N* and `isSamp`[*i*] = 0, this implies we do not know the position where *x* would occur in `useq`, and we must find the closest sampled position in order to decode the position of *x* (if it does, in fact, occur in `useq`). This is accomplished by Algorithm 1.

**Fig. 1A:**
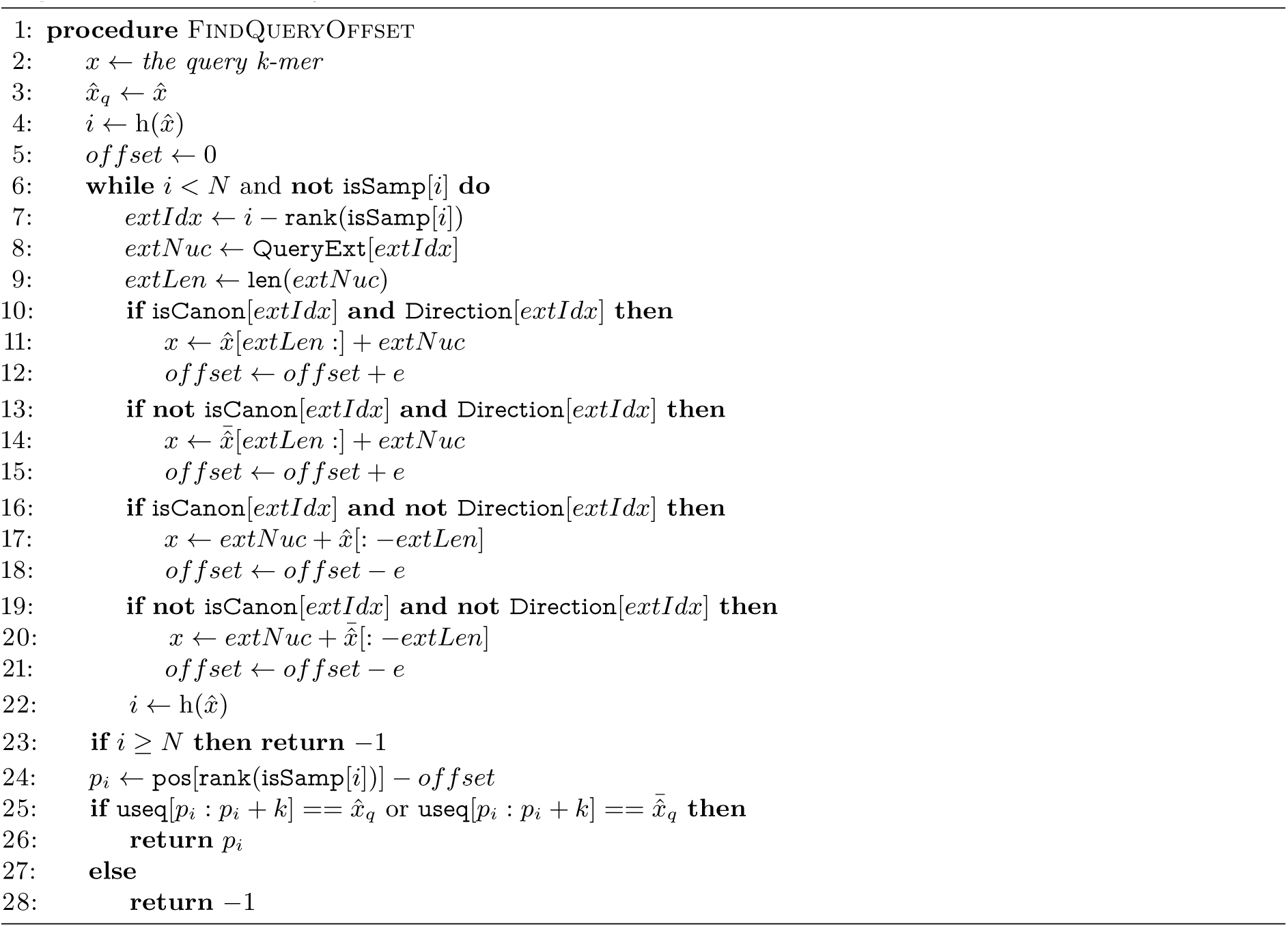
Algorithm 1. Find Query Offset

Intuitively, Algorithm 1 appends nucleotides stored in the `QueryExt` array to *x* to generate a new k-mer, *x*^′^, which either has a sampled position, or is closer to a sampled position than is *x*. The extension process is repeated with *x*^′^, *x*^″^, etc. until either an invalid position is returned by h, or a sampled position is reached. If an invalid position is returned at any point in the traversal, the original k-mer cannot have been a valid query. On the other hand, if a sampled position is reached, one still needs to verify that the k-mer implied by the query procedure is identical to the original k-mer query *x* (or 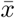). To check this, one simply traverses back to the position in `useq` for the original k-mer *x* that is implied by the sampled position and sequence of extension operations. The rest of the search proceeds as for the dense case. The whole process of a (successful) k-mer query in sparse index is illustrated in Figure 2 through an example.

**Fig. 2:**
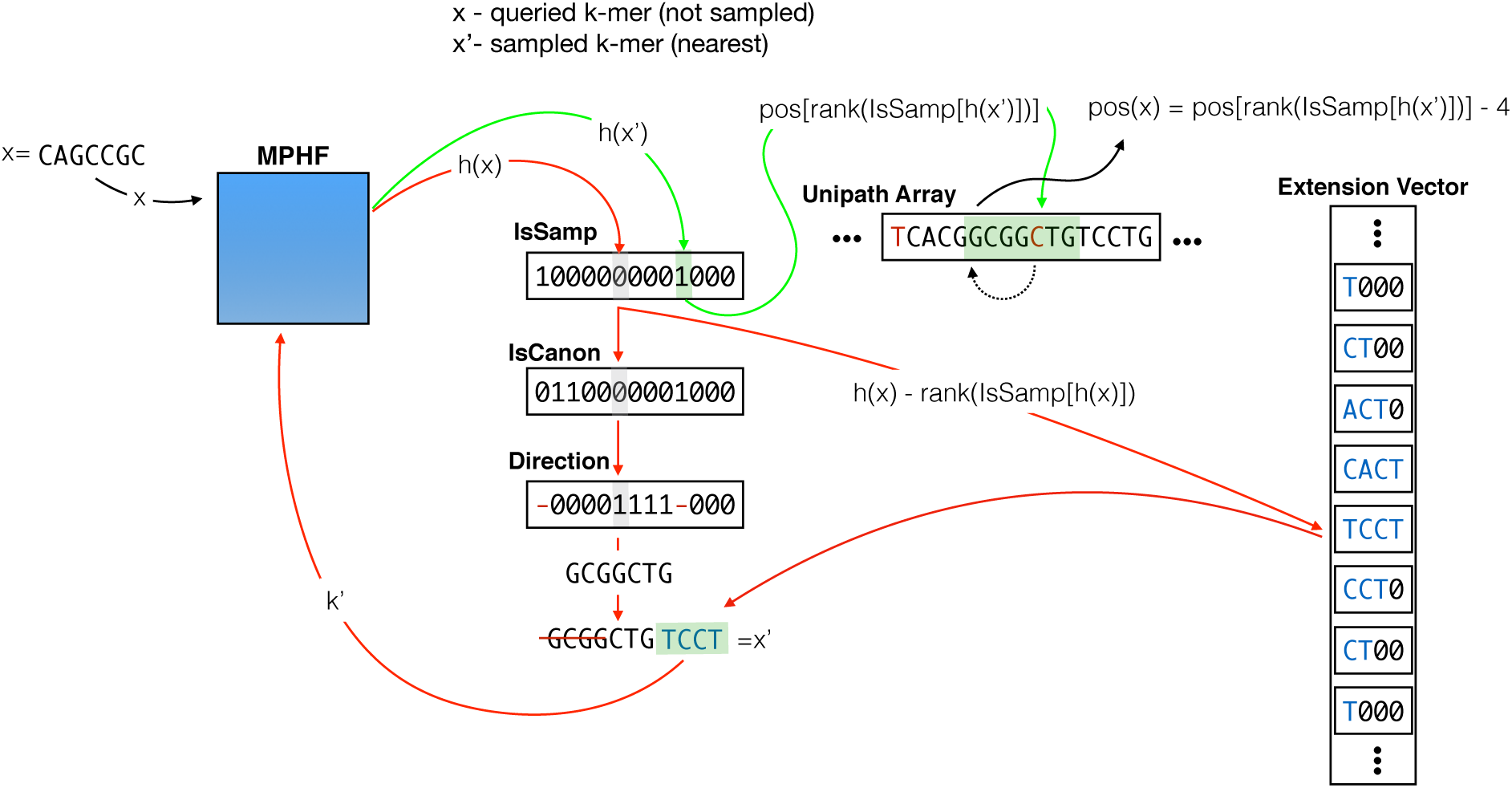
An illustration of searching for a particular k-mer in the *sparse* Pufferfish index with sample factor (*s*) of 9 and extension size (*e*) of 4. Vector isSamp has length equal to the number of valid k-mers, and `isCanon` and `Direction` have length equal to the total number of non-sampled k-mers. The minimum perfect hash yields the index 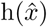 for *x* = `CAGCCGC` in `isSamp`, where we discover that the k-mer’s position is not sampled. Since `isCanon` 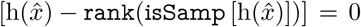 we know that the k-mer, if present, is not in the canonical orientation in `useq`. Since *x is* in the canonical orientation, we must reverse-complement it as 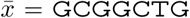 before adding the extension nucleotides. Then, based on the value of 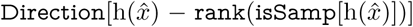, we know that to get to the closest sampled k-mer we need to append the extension nucleotides to the right of 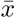 The extension is extracted from the `QueryExt` vector. Since extensions are recorded only for non-sampled k-mers, to find the index of the current k-mer’s extension, we need to determine the number of non-sampled k-mers preceding index 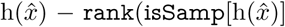. This can easily be computed as 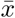, which is the index into `QueryExt` from which we retrieve this k-mers’s extension. We create a new k-mer, *x*^′^, by appending the new extension to 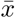, and also removing its first *e* = 4 bases. Then, we repeat the same process for the new k-mer *x*^′^. This time, the k-mer is sampled. Hence, we go directly to the index in `useq` suggested by pos 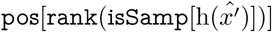. To check if the original k-mer we searched for exists, we need to compare the k-mer starting from *e* = 4 bases to the left of the current position with the *non-canonical* version of the original k-mer (since the sampled k-mer *x*^′^ was arrived at by extending the original query k-mer by 4 nucleotides to the right). Generally speaking, once we reach a sampled position, to check the original query k-mer, we need to move in `useq` to either the right or the left by exactly the distance we traversed to reach this sample, but in the opposite direction.

By altering the stored extension size *e* and the maximum sampling rate *s*, one can limit the maximum number of extension steps (and hence the maximum number of hash lookups) that must be performed in order to retrieve the potential index of *x* in `useq`. A denser sampling and longer extensions require fewer possible extension steps, while a sparser sampling and shorter extensions require less space for each non-sampled position. If 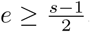, one can guarantee that at most a single extension step needs to be performed for any k-mer query, which allows k-mer queries to remain practically very fast while still reducing the index size for large reference sequences.

Even though the sparse index maintains a number of extra bit vectors not required by the dense index, it is usually considerably smaller. Assume a case where the extension length 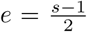 is approximately half of the sampling factor (the minimum length that will guarantee each query requires at most a single extension step). Since we keep the extension required to get to the closest position in the left or right direction, we need to keep *e* bases for a k-mer, with each base represented using 3 bits (since we need to allow encoding extensions of length *< e*, for which the encoding must allow a delimiter). Hence, this requires 3*e* bits per k-mer for the `QueryExt` vector. The `isCanon` and Direction vectors each require a single bit per non-sampled k-mer, and the `isSamp` vector requires a single bit for all *N* of the valid k-mers. Assume, for simplicity of analysis, that the sampled k-mers are perfectly evenly-spaced (which is not possible in practice since e.g., we must require to sample at least one k-mer from each unipath), so that the number of sampled k-mers is simply given by 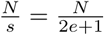. Further, since we are ignoring unipath boundary effects, assume that *N* = *L*_*s*_. Since the space required by the rest of the index components (e.g. the MPHF, and utab, etc.) is the same for the dense and sparse index, the sparse index will lead to a space savings whenever

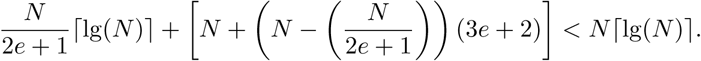

Under this analysis, in a typical dataset, such as the human genome with lg(*L*_*s*_) *≈* lg(*N*) *≈* lg(3 *×* 10^9^) *≥* 30 bits, and choosing *s* = 9 and *e* = 4, so that we sample every 9^th^ k-mer on average, and require at most one extension per query, we save, on average, *∼* 14.5 bits per k-mer. Of course, the practical savings are less because of the boundary effects we ignored in the above analysis.

## 3 Results

We explored the size of the index along with the memory and time requirements for index building and k-mer querying (a fundamental building block of many mapping and alignment algorithms) using Pufferfish and two other tools, BWA (BWA-MEM ^6^, specifically) and kallisto.

Though BWA is not a graph-based index, it was chosen as it implements the highly memory-efficient FMD-index ^6^, which is representative of a memory-frugal approach. It is also worth noting that, although we only test querying for fixed-length k-mers here, BWA is capable of searching for *arbitrary* length patterns — an operation not supported by the kallisto or Pufferfish indices. On the other hand, kallisto^162^ adopts a graph-based index, and provides very fast k-mer queries. Both BWA and kallisto implement all phases of index construction (i.e., the input to these tools is simply the FASTA files to be sequenced). For Pufferfish, however, we first need to build the compacted de Bruijn graph. We build the compacted de Bruijn graph and dump it in GFA format using TwoPaCo ^21^. Then (as the output does not satisfy our definition of a compacted de Bruijn graph) we need to further prepare the GFA file for indexing. We call this process *pufferization*. It converts the GFA file to the format accepted by Pufferfish (i.e., each k-mer should appear only once in either orientation among all the unipaths, and all unipaths connected in the compacted de Bruijn graph should have an overlap of exactly *k-*1 bases). Finally, we build both dense and sparse Pufferfish indexes and benchmark the time and memory for all steps of the pipeline individually. All experiments were performed on an Intel(R) Xeon(R) CPU (E5-2699 v4 @2.20GHz with 44 cores and 56MB L3 cache) with 512GB RAM and a 4TB TOSHIBA MG03ACA4 ATA HDD running ubuntu 16.10, and were carried out using a single thread except for compacted de Bruijn graph building step using TwoPaCo. For all datasets, we consider *k* = 31, and the sparse Pufferfish index was constructed with *s* = 9 and *e* = 4.

*References and query datasets* We performed benchmarking on three different reference datasets, selected to demonstrate how the different indices scale as the underlying reference size and complexity increases. Specifically, we have chosen a common human transcriptome (GENCODE version 25, 201 MB), a recent build of the human genome (GRCh38, 2.9 GB), and an ensemble of *>* 8000 bacterial genomes and contigs (18G) downloaded from RefSeq (ftp://ftp.ncbi.nlm.nih.gov/genomes/refseq/bacteria/). The human transcriptome represents a small reference sequence (which nonetheless exhibits considerable complexity due to e.g., alternative splicing), the human genome represents as a moderate (and very common) size reference, and the collection of bacterial genomes acts as a large reference set. For the k-mer query experiments, we search for all the k-mers from an experimental sequencing dataset associated with each reference. To query the human transcriptome, we use k-mers from SRA accession SRR1215997, with 10,683,470 reads, each of length 100 bases. To query the human genome, we use k-mers from SRA accession SRR5833294 with 34,129,891 reads, each of length 250 bases. Finally, to query the bacterial genomes, we use k-mers from SRA accession SRR5901135 (a sequencing run of *E. coli*) with 2,314,288 reads, each of length 250 bases.

*Construction time* The construction time for various methods depends, as expected, on the size and complexity of the references being indexed (Table 1). No tool exhibits faster index construction than all others across all datasets, and the difference in construction time between the fastest and slowest tools for any given dataset is less than a factor of 3. All tools perform similarly for the human transcriptome. For indexing the human genome, BWA is the fastest, follwed by Pufferfish and then kallisto. For constructing the index on all bacterial genomes, kallisto finished most quickly, followed by BWA and then Pufferfish. The time (and memory) bottleneck of index construction for Pufferfish is generally TwoPaCo’s construction of the compacted de Bruijn graph. This is particularly true for the bacterial genomes dataset where TwoPaCo’s compacted de Bruijn graph construction accounts for *∼*85% of the total index construction time. This motivates considering potential improvements to the TwoPaCo algorithm for large collections of genomes (as well as considering other tools which may be able to efficiently construct the required compacted de Bruijn graph input for Pufferfish).

**Table 1:** Upper half of the table shows construction time and memory requirements for BWA, kallisto and Pufferfish (dense and sparse) on three different datasets. In the lower half of the table, the construction statistics are provided for different phases of Pufferfish pipeline. The time requirement for Pufferfish is the sum of different sub parts of the workflow, where the memory requirement is the *max* of the same.

| Tool | Memory (MB) |  |  | Time (h:m:s) |  |  |
| --- | --- | --- | --- | --- | --- | --- |
|  | Human Transcriptome | Human Genome | Bacterial Genomes | Human Transcriptome | Human Genome | Bacterial Genomes |
| BWA | 292 | 4,443 | 32,213 | 2:56 | 0:58:27 | 13:11:45 |
| kallisto | 3,552 | 150,657 | 315,387 | 3:05 | 3:27:42 | 9:07:35 |
| pufferfish dense | 1,466 | 27,438 | 241,721 | 0:04:13 | 2:09:25 | 24:36:41 |
| pufferfish sparse | 1,466 | 27,438 | 241,721 | 0:04:41 | 2:28:53 | 25:12:52 |
| TwoPaCo | 1,466 | 18,004 | 241,721 | 0:02:47 | 0:56:12 | 21:25:46 |
| pufferize | 584 | 27,438 | 75,342 | 0:0:10 | 0:21:53 | 1:03:17 |
| pufferfish dense index | 438 | 20,000 | 50,459 | 0:01:16 | 0:51:20 | 2:07:38 |
| pufferfish sparse index | 331 | 17,745 | 50,457 | 0:01:44 | 1:10:48 | 2:43:49 |

## Construction memory usage

Unlike construction time, the memory required by the different tools for index construction follows a clear trend; BWA requires the least memory for index construction, followed by Pufferfish, and kallisto requires the most memory. There are also larger differences in the construction memory requirements than the construction time requirements. For example, to construct an index on the human genome, kallisto requires *∼* 34 times more memory than BWA (and *∼* 5.5 times more memory than Pufferfish). With respect to the current pipeline used by Pufferfish, we see that TwoPaCo is the memory bottleneck for the human transcriptome and bacterial genomes datasets, while *pufferize* consumes the most memory for the human genome. For the bacterial genomes dataset in particular, TwoPaCo consumes over 3 times as much memory as the next most intensive step (*pufferize*) and *∼* 4.8 times as much memory as actually indexing the input compacted de Bruijn graph. We note that TwoPaCo implements a multi-pass algorithm, which can help control the peak memory requirements in exchange for performing more passes (and therefore taking longer to finish). However, we did not thoroughly explore different parameters for TwoPaCo’s Bloom filter size (which indirectly affects the number of passes).

*Query time and memory* To measure the query time required for k-mer lookup by the different methods, we performed experiments in which k-mer queries were issued from sequencing read sets related to each of the reference sequences (see the description of the query datasets above). Specifically, for each method and each dataset, we measure the time it takes to load the index into memory, stream through each valid k-mer in each read (i.e., k-mers not containing non-ATCG characters), and record the total number of occurences of all queried k-mers in the corresponding reference index. By querying with experimental data, we mimic a realistic distribution of queries for both present and absent k-mers in the index. Since we only perform k-mer queries here (and not read mapping or alignment), we consider only the first end of paired-end read datasets. Comparing the times recorded in Table 2, we can see that Pufferfish (both the dense and sparse variants) and kallisto generally tend to complete the query task faster than BWA, except in the bacterial genomes dataset, where BWA takes less time than kallisto (partly because of the time required by kallisto to load its large index on this dataset). Again, this can be attributed, at least in part, to the fact that both Pufferfish and kallisto are k-mer-based indices, while BWA provides a full-text index. However, much of the speed of the hashing-based solutions may also be attributed to the efficiency of hashing as a lookup scheme and to the ability of the compacted de Bruijn graph to efficiently represent highly-repeated patterns. As expected, query in the sparse Pufferfish index is slower than in the dense index, though the sparse index still remains practically fast in these benchmarks.

When examining the memory required for querying, we observe a pattern similar to that which we saw with construction memory. That is, BWA requires the least memory, followed by Pufferfish, and kallisto requires the most memory. Here, however, the gap between BWA and Pufferfish is reduced, as the final Pufferfish index typically consumes considerably less memory than is required during construction (especially for the bacterial genomes dataset). In fact, we see that the disk space and query memory requirements of Pufferfish are very similar, as is the case with BWA. For kallisto, however, the hash table consumes much more memory than does the serialized index on disk^*†*^. The difference in memory requirements is particularly striking for the large and diverse bacterial genomes dataset.

**Table 3:**
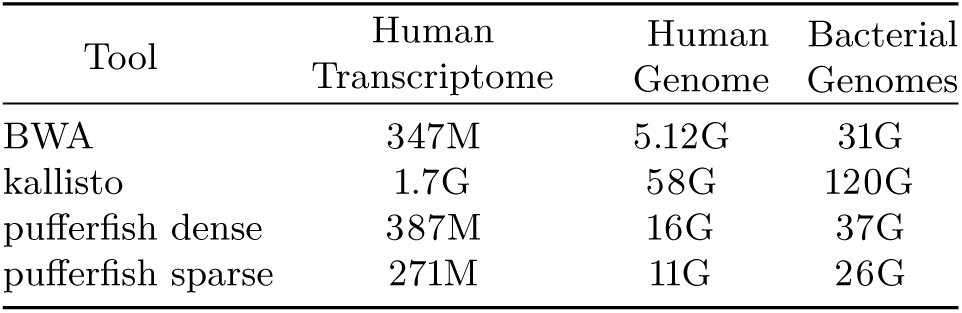
Disk space required for the index of each tool on different datasets.

## 4 Conclusion and Future Work

In this paper we proposed a new efficient data structure for indexing compacted colored de Bruijn graphs, and implement this data structure in a tool called Pufferfish. We showed how Pufferfish can achieve a balance between time and space resources. By building upon a MPHF ^27^, we provide practically fast k-mer lookup, and by carefully organizing our data structure and making use of succinct representations where applicable, we greatly reduce the space compared to traditional hashing-based implementations. The main components of the data structures are a minimum perfect hash function (MPHF) built on k-mers, the concatenated unipath array from which the k-mers are sampled, a bit vector that marks the boundary of unitigs in the concatenated array, a vector containing the offset position for the k-mers, and a unipath table enumerating the occurrences of each unipath in the reference sequences.

Moreover, we presented two variants of the Pufferfish data structure; namely, a dense and a sparse variant. The first is optimized for fast queries and the second provides the user with the ability to trade off space for speed in a fine-grained manner. In the sparse index, we only keep offset positions for a subset of k-mers. To query a k-mer whose position is not sampled, the sparse representation is aided with a few auxiliary data structures of much smaller size. Since the largest component of the index is the position vector, adopting this sparse representation significantly reduces the required memory and disk space. Our analyses suggest that Pufferfish (dense) achieves similar speed to existing hash-based approaches, while greatly reducing the memory and disk space required for indexing, and that Pufferfish (sparse) reduces the required space even further, while still providing fast query capabilities. We consider indexing and query on both small (human transcriptome) and large (*>* 8000 bacterial genomes) reference datasets. Pufferfish strikes a desirable balance between speed and space usage, and allows for fast search on large reference sequences, using moderate memory resources.

Having built an index for a reference genome, transcriptome, or metagenome using Pufferfish, the immediate future work consists of implementing relevant applications based on this index. These applications fall into the categories of problems that need mapping or alignment as their initial step. Therefore, we would like to build a fast mapper or aligner by adopting an algorithm such as that described in Sarkar et al.^29^, and modifying it to use Pufferfish as its indexing methodology. Later, we can use the result of this aligner for mapping to a population of genomes, and performing downstream tasks such as contaminant detection, metagenomic abundance estimation, etc. We expect the memory efficiency of Pufferfish will be beneficial in working with larger collections of genomic and transcriptomic and metagenomic datasets.

## Footnotes

* https://github.com/GFA-spec/GFA-spec

** Specifically, the TwoPaCo compacted de Bruijn graph has two main differences with the format that is expected by Pufferfish. First, it is not the case that k-mers and their reverse complements will appear only once in the TwoPaCo compacted de Bruijn graph. Second, the GFA generated by TwoPaCo assumes that *edges* of size at least *k* + 1 will act as GFA segments, implying that they will overlap by *k* nucleotides. However, we require that segments be of at least size *k* and overlap by exactly *k-*1 nucleotides.

* * * Throughout this manuscript, we adopt Python notation to represent string manipulation, and string slices, prefixes and suffixes.

† We also note that on large sequences (e.g., the human genome and bacterial genomes), kallisto seemed to require an inordinate amount of time (i.e., days) to load the index into memory. This occurred during the final phase of index loading. However, by modifying the index loading code, we were able to resolve this issue and hence provide k-mer query times for these samples.

